# SECANT: a biology-guided semi-supervised method for clustering, classification, and annotation of single-cell multi-omics

**DOI:** 10.1101/2020.11.06.371849

**Authors:** Xinjun Wang, Zhongli Xu, Xueping Zhou, Yanfu Zhang, Heng Huang, Ying Ding, Richard H. Duerr, Wei Chen

## Abstract

The recent advance of single cell sequencing (scRNA-seq) technology such as Cellular Indexing of Transcriptomes and Epitopes by Sequencing (CITE-seq) allows researchers to quantify cell surface protein abundance and RNA expression simultaneously at single cell resolution. Although CITE-seq and other similar technologies have quickly gained enormous popularity, novel methods for analyzing this new type of single cell multi-omics data are still in urgent need. A limited number of available tools utilize data-driven approach, which may undermine the biological importance of surface protein data. In this study, we developed SECANT, a biology-guided SEmi-supervised method for Clustering, classification, and ANnoTation of single-cell multi-omics. SECANT can be used to analyze CITE-seq data, or jointly analyze CITE-seq and scRNA-seq data. The novelties of SECANT include 1) using confident cell type labels identified from surface protein data as guidance for cell clustering, 2) providing general annotation of confident cell types for each cell cluster, 3) fully utilizing cells with uncertain or missing cell type labels to increase performance, and 4) accurate prediction of confident cell types identified from surface protein data for scRNA-seq data. Besides, as a model-based approach, SECANT can quantify the uncertainty of the results, and our framework can be easily extended to handle other types of multi-omics data. We successfully demonstrated the validity and advantages of SECANT via simulation studies and analysis of public and in-house real datasets. We believe this new method will greatly help researchers characterize novel cell types and make new biological discoveries using single cell multi-omics data.

## Introduction

Single-cell RNA-sequencing (scRNA-seq) technologies have advanced rapidly for understanding cell heterogeneity and discovering rare cell types from normal and disease tissues (Treutlein et al. 2014; Grun et al. 2015; Gawad et al. 2016; Tsoucas and Yuan 2017; Yuan et al. 2017). Embedded in the popular scRNA-seq platform such as 10X Genomics Chromium System (Zheng et al. 2017), the recently developed CITE-seq (Cellular Indexing of Transcriptomes and Epitopes by sequencing) (Stoeckius et al. 2017a) (or similar REAP-seq (RNA expression and protein sequencing) (Peterson et al. 2017)), and cell hashing technologies (Stoeckius et al. 2018) allow for immunophenotyping of single cells based on cell surface expression of specific proteins together with simultaneous transcriptome profiling and sample origin detection within a cell. Besides, more omics types of single cell data are emerging (Buenrostro et al. 2015; Cusanovich et al. 2015; Mimitou et al. 2020). In these single cell multi-omics experiments, the abundance of different kinds of features such as mRNA or cell surface protein is converted into a quantitative and sequenceable readout through the use of DNA-barcoded antibodies and can be measured by the count of Unique Molecular Index (UMI) and Antibody-Derived Tags (ADT), respectively, simultaneously at single cell resolution.

Although there are a large number of existing tools for analyzing droplet-based scRNA-seq data (Satija et al. 2015; Ji and Ji 2016; Kiselev et al. 2017; Lopez et al. 2018; Sun et al. 2018; Wang et al. 2018; Sun et al. 2019), model-based statistical methods for analyzing single-cell multi-omics data are still in urgent need. We will focus on analyzing CITE-seq data, one of the most informative multi-omics types, in this paper although the method can be generalized to other type of data. For convenience, we will use ADT data to denote surface protein data in this paper. CITE-seq refers to single cell multi-omics data with both scRNA-seq and ADT measurements for each cell. There are a number of cutting-edge methods for surface protein imputation with scRNA-seq data (Gayoso et al. 2020; Zhou et al. 2020) and for joint clustering of both protein and RNA features (Gayoso et al. 2020; Wang et al. 2020). These methods, although having demonstrated their outstanding performance, all tend to utilize a data-driven approach but do not use much of existing biological knowledge. For example, it is a common approach in joint clustering methods to integrate protein and RNA data by transforming both features onto a similar space. However, by doing so, the important underlying biological information from surface protein marker could be undermined. On the contrary, biological researchers often consider the cell surface markers as gold standard to define cell types in molecular biology, where researchers identify distinguished cell types through cell gating such as flow cytometry with a list of classic differentiation (CD) markers, such as CD3, CD4, CD8 and CD19 (Maecker et al. 2012; Verschoor et al. 2015; Lian et al. 2020). Thus, a more biological knowledge driven approach should consider to give more weight on ADT data for the purpose of cell clustering and cell type identification. For example, use ADT data to first label the well-defined (confident) cell types, such as B cells, Monocytes, CD4+ T cells, CD8+ T cells and natural killer (NK) cells, and then use this information as a guidance for clustering with RNA data. This approach utilizes a great amount of biological knowledge to avoid a common issue that some identified clusters are mixtures of multiple general cell types, for example, CD4+ T cells and CD8+ T cells, from clustering with RNA data (Chen et al. 2002; Haider and Pal 2013; Stoeckius et al. 2017b). Despite cell clustering, ADT data can also play a role in cluster annotation. Current methods for cluster annotation rely on post-hoc differential expression (DE) analysis on RNA data, and researchers need to select a couple of plausible DE gene markers from a long list. However, it is often challenging since gene markers are not always correlated with their corresponding surface markers. Thus, we expect it vastly beneficial to provide some confident cell type annotation from ADT data for researchers to further figure out the identities of cell subtypes. Another popular research topic is to jointly analyze data from CITE-seq and scRNA-seq, by which we can assume the cell compositions are similar though batch effect exists. There are many advantages of joint analyzing CITE-seq and scRNA-seq data, e.g., the addition of an extra RNA data could help increase clustering performance due to larger sample size, and we can also provide the confidence cell type annotation from ADT data to scRNA-seq data.

Motivated by the above demands, in this study we propose a novel framework for surface protein guided cell clustering and general cluster annotation with CITE-seq data. If more scRNA-seq datasets from similar cell populations are available, SECANT can be used to jointly analyze data from CITE-seq and scRNA-seq to predict confident cell types for scRNA-seq data, and enhance the performance of cell clustering and cluster annotation. Our method utilizes a model-based approach and is developed based on several statistical models in semi-supervised learning (Bouveyron et al. 2019). As a biological knowledge-driven approach, the input of our method from ADT data is the confident cell type label, which can be obtained using cell gating or other existing methods. To overcome a common issue that some cells are hard to identify cell types even with ADT data, instead of excluding those cells from the analysis, which will cause the loss of sample size and potential the drop of some novel cell subtypes, our method is specifically designed to accommodate those cells with “uncertain” labels into our model so that we can fully utilize their transcriptomic information. We use extensive simulation studies to demonstrate the validity of our proposed method, and we illustrate the usefulness and easy interpretation of our method with two real data applications.

## Results

### General workflow of SECANT

The general workflow of SECANT is shown in Figure 1. Our method can work with CITE-seq data (scRNA + ADT) only or jointly analyze CITE-seq and scRNA-seq data. For using CITE-seq data only, the raw data matrices need to first undergo some data pre-processing steps, and the inputs of our method include the confident cell type labels built from ADT data and the latent space of RNA data after dimension reduction. Our method considers ADT cell type labels as general guidance for cell clustering with RNA data by introducing certain constraints through a probabilistic concordance matrix. We establish a statistical model and maximize the log-likelihood of the observed data to estimate the concordance matrix and ADT-guided cell clustering results. Through the estimated concordance matrix, we can provide confident cell type annotation for each cluster, for example cluster 1 and cluster 2 are potential sub-clusters of B cells. For joint CITE-seq and scRNA-seq analysis, we require that the latent spaces of pooled RNA data are similar in distribution so the clustering parameters are commonly shared by both data. The inclusion of the RNA data from scRNA-seq into the model could increase the precision of estimating concordance matrix as well as ADT-guided cell clustering results. Also, our model can predict the ADT cell type labels for cells from scRNA-seq experiment, which does not have ADT information. Other important benefits of our method include utilizing cells with uncertain cell type label from ADT data, and providing uncertainty of the results (“soft labels”) in terms of posterior probability. Details about our statistical models can be found in Methods section.

**Figure 1.**
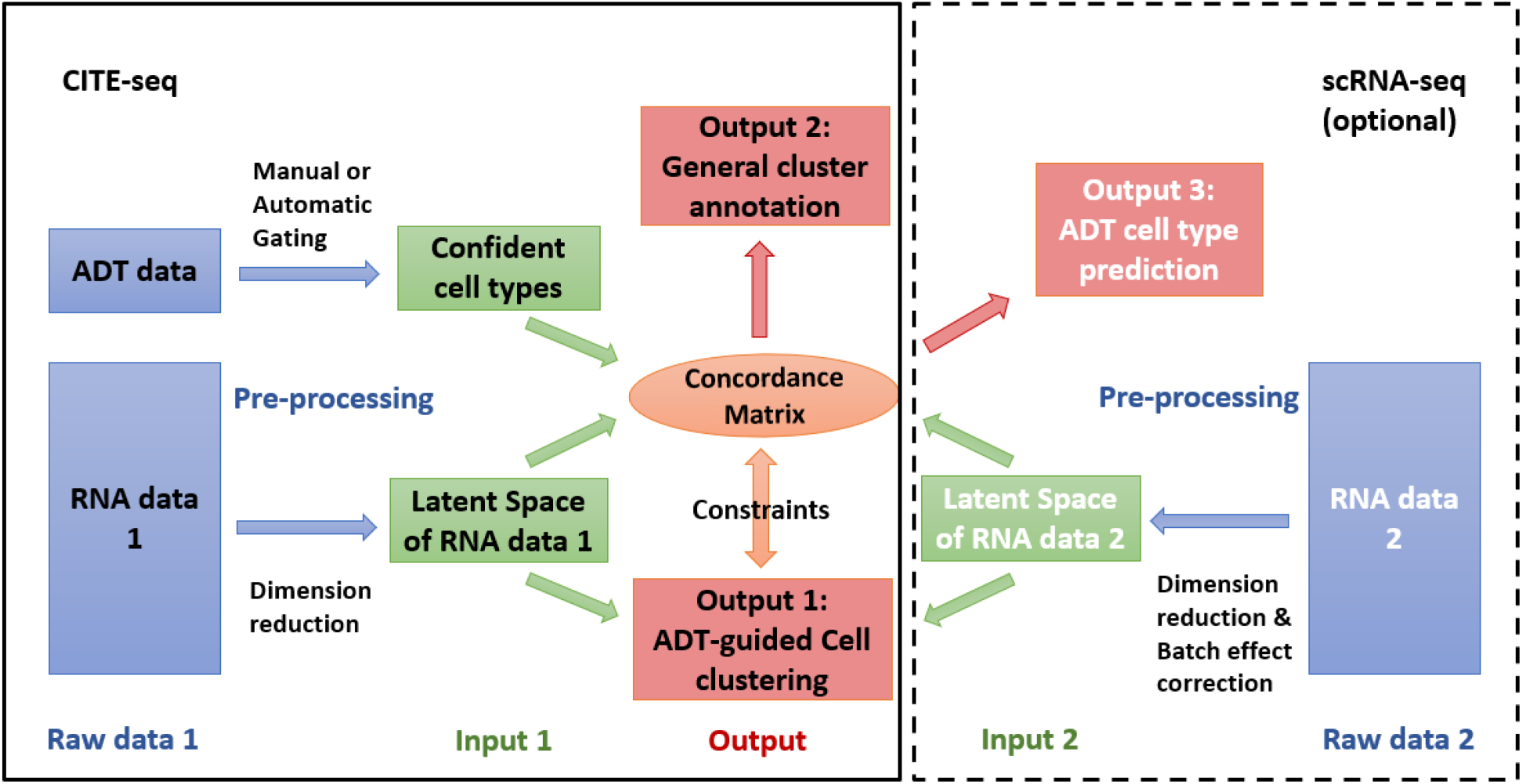
General workflow of SECANT. In this study, we used manual gating to classify confident cell types with ADT data, and used scVI for dimensional reduction and batch effect correction with RNA data for data pre-processing.

### Identifying confident cell types with ADT data

In the first step, we obtain the confident cell type label directly through manual gating with ADT surface marker data, motivated by the fact that it has been a widely used tool for cell type identification with flow cytometry and mass cytometry data, and the corresponding pipelines proposed by the biologists are quite mature (Maecker et al. 2012; Verschoor et al. 2015). In addition, the bi-modal or multi-modal mixture Gaussian distribution structure of log-transformed ADT count fits gating pipeline naturally. In Supplemental Figure 1, we summarize a workflow to illustrate how to gate some general cell types for peripheral blood mononuclear cells (PBMC) from ADT data. However, one of the major challenges of manual gating is its subjective choice of gating boundary. In general, a less stringent boundary will introduce mixture of target cells with other cell types, while a more stringent boundary will lead to less target cells to be identified (Supplemental Fig. 2). To overcome this challenge, our method is designed to utilize cells with uncertain cell type labels. It is worth noting that tools other than manual gating can also be used for identifying confident cell types with ADT data (Lian et al. 2020).

### Pre-processing of scRNA-seq data

RNA data need to undergo pre-processing for dimension reduction and batch effect correction. In this study, we apply scVI for both purposes. scVI is a popular tool that utilizes variational autoencoder for non-linear dimensional reduction and batch effect correction (Lopez et al. 2018). We also tested the performance of batch effect correction on two public PBMC datasets (Supplemental Fig. 3). To be specific, we first processed RNA data with scVI, and the resulting latent space, which follows a low-dimensional Gaussian distribution, will then be used as the input of our method. In general, other tools for dimension reduction and batch effect correction can also be used for data pre-processing in our proposed framework, although the distribution assumed in the statistical model is subject to change for better data fit.

### Simulation results

We first assessed the performance of SECANT with simulation studies. For convenience, we simulated data from mixture multivariate Gaussian distribution to mimic the latent space of RNA data after applying scVI. For general setting, we simulate 8 cell clusters and label them with 4 confident cell types, which are used to mimic the input from ADT data. The distribution parameters used to generate data are decided based on the estimates obtained from applying SECANT on a public PBMC dataset. Further, to mimic the practical situation that we usually fail to identify the cell type for some cells, we randomly assign a certain proportion, denoted by *p^U^*, of cells with uncertain label. We simulated 100 datasets under each simulation setting. In Figure 2 we use Uniform Manifold Approximation and Projection (UMAP) plot (McInnes et al. 2018), a popular non-linear dimension reduction tool used in single cell analysis, to visualize an example of our simulated data (2000 cells, 20% with uncertain label). Figure 2A is colored by the 4 confident cell types (type 1 - type 4), which refer to B cells, CD14+ Monocytes, CD4+ T cells and CD8+ T cells, respectively. Figure 2B is colored by the 8 clusters (cluster 1 - cluster 8), where cluster 1 belongs cell type 1, cluster 2 and 3 belong to cell type 2, cluster 4 and 5 belong to cell type 3, and cluster 6, 7 and 8 belong to cell type 4. This simulation setting could largely reflect the real-world scenario that CD4+ T cells and CD8+ T cells are often clustered together with scRNA-seq data.

**Figure 2.**
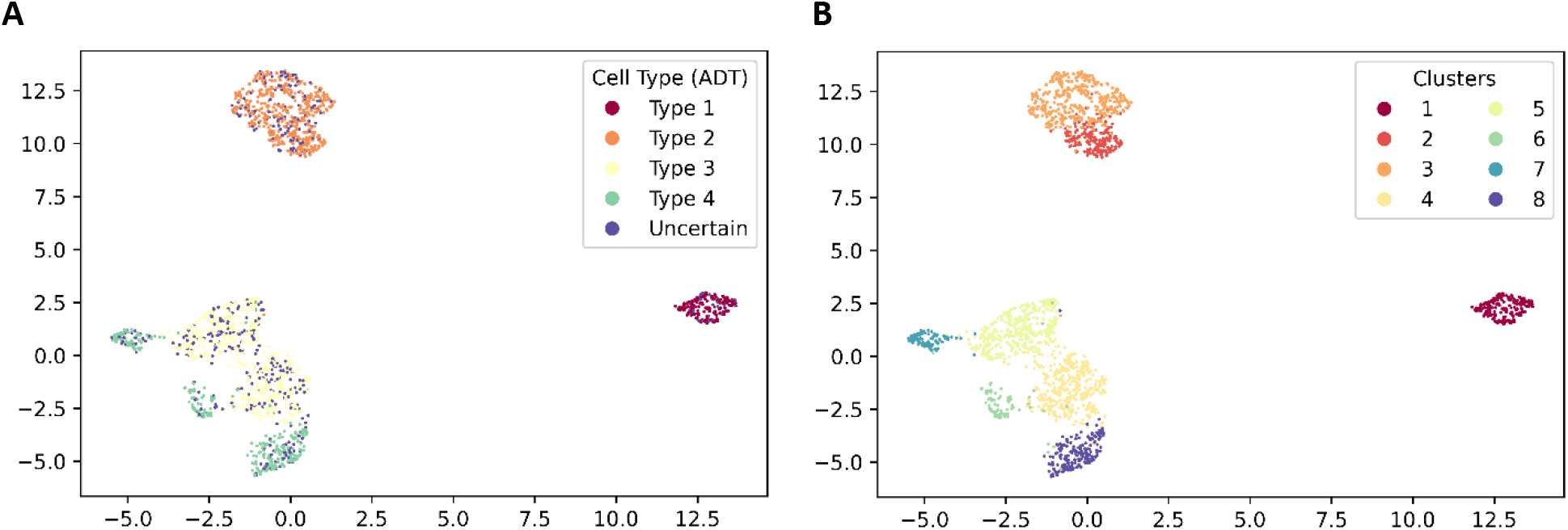
UMAP of an example simulated data under our simulation setting. 2A is colored by simulated true cell types (mimic input from ADT data). 2B is colored by the simulated true cluster assignments.

### Performance of ADT-guided clustering with one data input

We assessed the performance of ADT-guided clustering of SECANT by computing adjusted random index (ARI) (Rand 1971) and adjusted mutual information (AMI) (Nguyen et al. 2009) with the simulated clustering truth. Both ARI and AMI are commonly used metrics for the concordance of two clustering results. Comparing to the truth, an ARI or AMI of value 1 indicates the clustering result is identical to the truth, while value 0 indicates the clustering result is a random assignment. A previous study suggests using ARI for balanced clustering situation, while using AMI for unbalanced clustering situation (Romano et al. 2016). We also applied K-means and multivariate Gaussian mixture model (GMM) on the simulated data and summarized their results for reference. We alter the total number of cells, denoted by *N*, as well as the proportion of cells randomly assigned with the uncertain label, *p^U^*. In Figure 3A, we show the boxplot of ARI from SECANT under different *p^U^* settings, as well as from K-means and multivariate GMM, and the total number of cells varies from 500 to 2000. We observe that the clustering performance of SECANT increases with larger sample size, but decreases with larger *p^U^*. Indeed, we get less information from ADT cell type label with larger *p^U^*. On the other hand, although the performance of multivariate GMM and K-means also increases with larger sample size, our proposed SECANT performs the best among the three under all scenarios, even when 60% of cells are labeled as “uncertain” in the input ADT cell type label. The results of AMI show consistent pattern as ARI (Supplemental Fig. 4A).

**Figure 3.**
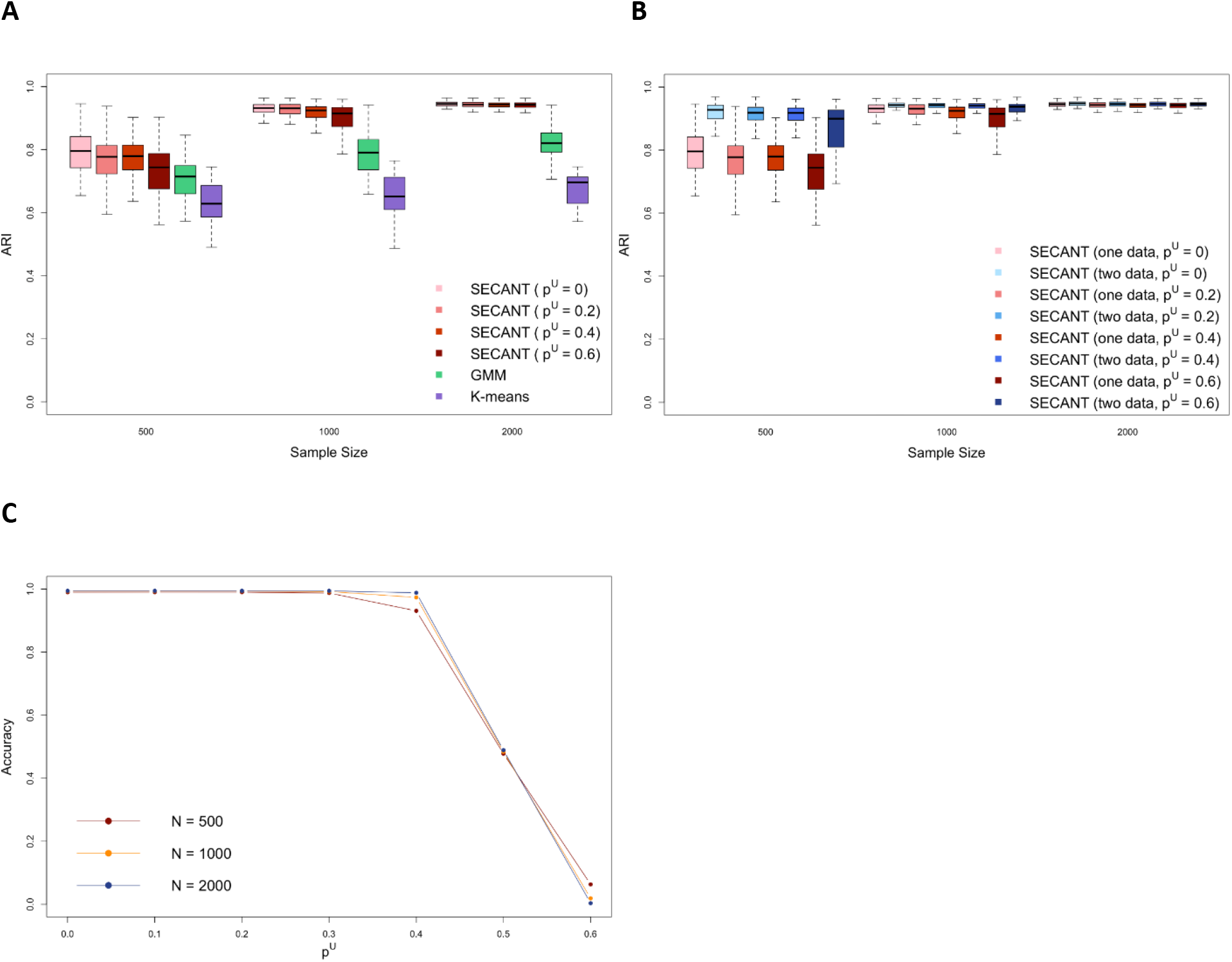
Results from simulation studies. 3A shows the distribution of ARI of SECANT (with different *p^U^* setting), K-means and GMM across different sample size *N*. 3B shows the distribution of ARI of SECANT using one dataset input vs. two datasets input under various *p^U^* and *N* settings. 3C shows the trend of ADT cell type label prediction accuracy with various *p^U^* and *N* settings.

### Performance of joint analysis with two data input

To assess the performance of SECANT for joint analysis of CITE-seq and scRNA-seq data, we first generated two datasets from the same distribution for each simulation, and the data generation method is the same as described above. Therefore, both datasets are composed of ADT label and RNA data. Next, we masked the ADT label from one dataset to mimic scRNA-seq data. Under this simulation setting, we will evaluate if the additional “unlabeled” scRNA-seq data could help increase the clustering performance, and assess the prediction accuracy of ADT cell type label for the dataset whose label is masked. In Figure 3B, we compare the clustering performance (ARI) of SECANT using CITE-seq data only and using both CITE-seq and scRNA-seq data. The sample size *N* for each dataset varies from 500 to 2000, and *p^U^* varies from 0 to 0.6. In addition to the similar patterns in Figure 3A, we find that the inclusion of an additional scRNA-seq data into our model can help increase the clustering performance, especially when sample size is small. The results of AMI show consistent pattern as ARI (Supplemental Fig. 4B). As for prediction accuracy, the results are summarized in Figure 3C. Although it is observed that the mean predication accuracy decreases with increasing *p^U^*, the accuracy actually remains very high if the uncertain rate of input label is less than 40%. The performance breaks down drastically when *p^U^* is greater than 50%, which is as expected since more than half of the input label provides no information. Similar results are found in robust mixture discriminant analysis (RMDA) (Bouveyron and Girard 2009). Also, it is interesting to observe that the prediction accuracy increases with larger sample size when *p^U^* is smaller than 50%, but such a trend is reversed when *p^U^* is greater than 50%. We further evaluated the cause of the resulting low prediction accuracy when *p^U^* is greater than 50%, and found that most cells are predicted as “uncertain” by our model when *p^U^* is high (Supplemental Figure 4C). Overall, we demonstrate the validity and the outstanding performance of SECANT for analyzing CITE-seq data only as well as joint analysis of CITE-seq and scRNA-seq data though simulation studies.

### Performance of detecting novel cell cluster

In practice, there could exist some novel cell clusters that we fail to identify their confident cell type label from ADT data due to lack of biological knowledge. For example, there are always some residual cells from cell gating, and those cells are classified as “uncertain” type in our analysis. To mimic this real-world scenario, we changed the confident cell type label of cells in cluster 8 from “Type 4” (referring to CD8+ T cells) to “uncertain” to pretend we don’t have enough knowledge to identify the cell type for those cells (Supplemental Fig. 5A). The clustering results under the guidance of new ADT label are shown in Supplemental Fig. 5B, and we observe that SECANT successfully identify this “novel” cluster as cluster 6, which is not mixed by nearby clusters (ARI=0.949; AMI=0.946). As for cell type annotation, which can be concluded from the estimated concordance matrix (Supplemental Table 1), we observe that cluster 6 is labeled as 100% “uncertain”, indicating this cluster is not the subtype of any existing confident cell types identified from ADT data. The results are consistent with our simulation setting.

### Joint analysis of two public PBMC CITE-seq datasets

We downloaded two public PBMC CITE-seq datasets from 10X Genomics website, named 10X10k_PBMC (7865 cells) and 10X5k_PBMC (5527 cells). For each dataset, we obtained 5 confident cell type labels, including B cells, CD14+ Monocytes, CD4+ T cells, CD8+ T cells, and Natural Killer (NK) cells, from ADT data using the gating procedure as illustrated in Supplemental Figure 1. Note that we didn’t assign a specific cell type label for CD16+ Monocytes due to its low amount, and those corresponding cells are classified as “uncertain”. The proportion of cells classified in the “uncertain” group is 16.3% for 10X10k_PBMC dataset, and 11.5% for 10X5k_PBMC dataset. For both RNA data, we applied scVI for batch effect correction and dimension reduction, and the latent space is 10-dimensional. For the purpose of joint analysis with paired CITE-seq data and scRNA-seq data, we temporarily removed the ADT data from 10X5k PBMC dataset and used only its RNA data together with 10X10k_PBMC CITE-seq data as the input.

We set the total number of clusters to be 11, and restrict the maximum number of clusters a confident cell type can have to be 3, which results in a total number of 45 different configurations of matrix form of the concordance matrix. We ran our method in parallel and decided the best matrix form based on log-likelihood. According to the best matrix form, there are 2 cell clusters corresponding to B cells, 2 corresponding to CD14+ Monocytes, 3 corresponding to CD4+ T cells, 3 corresponding to CD8+ T cells, and 1 corresponding to NK cells. In Figure 4, we visualize the ADT-guided clustering results for 10X10k_PBMC dataset. The UMAP plots are constructed based on the latent space of RNA data from 10X10k_PBMC dataset. Figure 4A is colored by the confident cell types built with ADT data, and Figure 4B is colored by the ADT-guided clustering results. We observe that none of the clusters is obviously a mixture of multiple ADT confident cell types. Similar results are also observed for 10X5k_PBMC dataset (Supplemental Fig. 7A-7B). As a comparison, we also ran multivariate GMM on the latent space of RNA data from 10X10k_PBMC dataset, and ran Seurat (Satija et al. 2015) with raw RNA data matrix, and the results are shown in Supplemental Figure 6. For multivariate GMM, we observe that cluster 3 is a mixture of NK cells and CD8+ T cells, and cluster 9 is a mixture of CD4+ T cells and CD8+ T cells. For Seurat, cluster 4 is a mixture of CD4+ T cells and CD8+ T cells, although the majority are CD8+ T cells. Similar results are also observed for 10X5k_PBMC dataset (Supplemental Fig. 7C-7D), and the mixture cluster issue tends to be worse, possibility due to a smaller sample size. Thus, we demonstrate the validity of our method. The estimated concordance matrix is shown in Table 1A, and we observe that the proportion of uncertain cells in cluster 3 and cluster 6 is more than 50%, which indicate that those clusters could be novel clusters that don’t belong to any common cell types. Based on the estimated concordance matrix, we can provide confident cell type annotation for the identified 11 clusters. The results can be visualized through an alluvial plot (Figure 4C). We further did post-hoc DE analysis based on gene markers, and successfully identify 9 out of 11 clusters based on common biological knowledge and existing literatures (show in Table 1B) (Choi et al. 2015; Martin and Badovinac 2018; Sampath et al. 2018).

**Figure 4.**
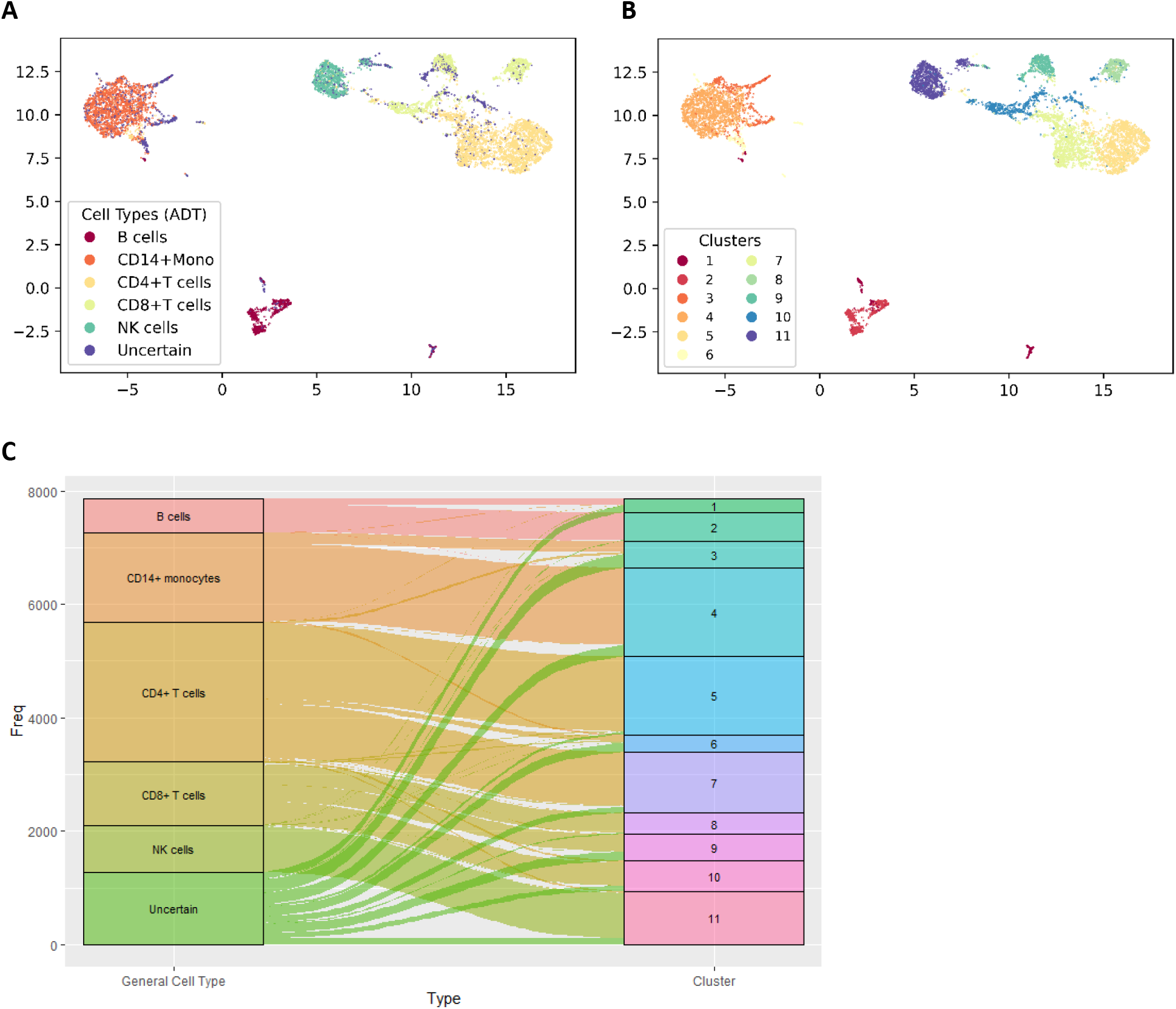
ADT-guided clustering results for 10X10k_PBMC data. UMAP plot is constructed using the latent space of RNA data. 4A is colored by confident cell types identified with ADT data. 4B is colored by the ADT-guided clustering results by SECANT. 4C shows the alluvial plot indicating the correspondence between confident cell type (provided by ADT data) and cell clusters.

**Table 1.** Concordance matrix (1A) and post-hoc subtype identification (1B) for public PBMC datasets.

**A**
|  | 1 | 2 | 3 | 4 | 5 | 6 | 7 | 8 | 9 | 10 | 11 |
| --- | --- | --- | --- | --- | --- | --- | --- | --- | --- | --- | --- |
| B cells | 0.56 | 0.97 | 0 | 0 | 0 | 0 | 0 | 0 | 0 | 0 | 0 |
| CD14+ Monocytes | 0 | 0 | 0.45 | 0.89 | 0 | 0 | 0 | 0 | 0 | 0 | 0 |
| CD4+ T cells | 0 | 0 | 0 | 0 | 0.98 | 0.46 | 0.91 | 0 | 0 | 0 | 0 |
| CD8+ T cells | 0 | 0 | 0 | 0 | 0 | 0 | 0 | 0.95 | 0.66 | 0.77 | 0 |
| NK cells | 0 | 0 | 0 | 0 | 0 | 0 | 0 | 0 | 0 | 0 | 0.89 |
| Uncertain | 0.44 | 0.03 | 0.55 | 0.11 | 0.02 | 0.54 | 0.09 | 0.05 | 0.34 | 0.23 | 0.11 |
| Cluster proportion<br>(10X10k_PBMC) | 2% | 6% | 8% | 19% | 24% | 5% | 13% | 4% | 4% | 7% | 10% |

**B**
|  | DE markers | Subtype Identification |
| --- | --- | --- |
| Cluster 1 |  | Marginal zone B cells |
| Cluster 2 | MZB1 | Follicular B cells |
| Cluster 3 |  | Intermediate Monocytes* |
| Cluster 4 | CLEC10A | Classical Monocytes |
| Cluster 5 | LEF1; TCF7 | T follicular helper cells* |
| Cluster 6 |  | Unknown |
| Cluster 7 | GATA3 | Th2 cells |
| Cluster 8 | CCR7 | Central memory CD8+ T cells* |
| Cluster 9 |  | Unknown |
| Cluster 10 | GZMK, GZMM | Cytotoxic T cells |
| Cluster 11 |  | NK cells |

Note: Identifying cell types labeled with \* are supported by existing literatures.

To assess the performance of ADT cell type prediction, we compared the predicted cell type labels with the actual confident cell type labels built from 10X5k_PBMC ADT data, which was previously masked when running our algorithm. The results are visualized in Figure 5, and we observe that the major differences are among those cells that are classified as “uncertain” or predicted as “uncertain”. In general, the predicted labels are quite close to actual labels. We also computed the confusion matrix between the predicted general cell types and the actual confident cell types (Supplemental Table 2). Excluding cells that are either classified as “uncertain” or predicted as “uncertain”, the overall prediction accuracy achieves 94.5%. Since our method is based on a probabilistic model, we can provide uncertainty of our results. In Supplemental Figure 8, we compared the distribution of maximum posterior probability for cells whose labels are correctly predicted and for cells whose labels are incorrectly predicted, and we observe the former cells in general have a much higher posterior probability compared to latter cells, indicating we are much more confident about the predicted cell labels for cells we indeed correctly predicted their confident cell types.

**Figure 5.**
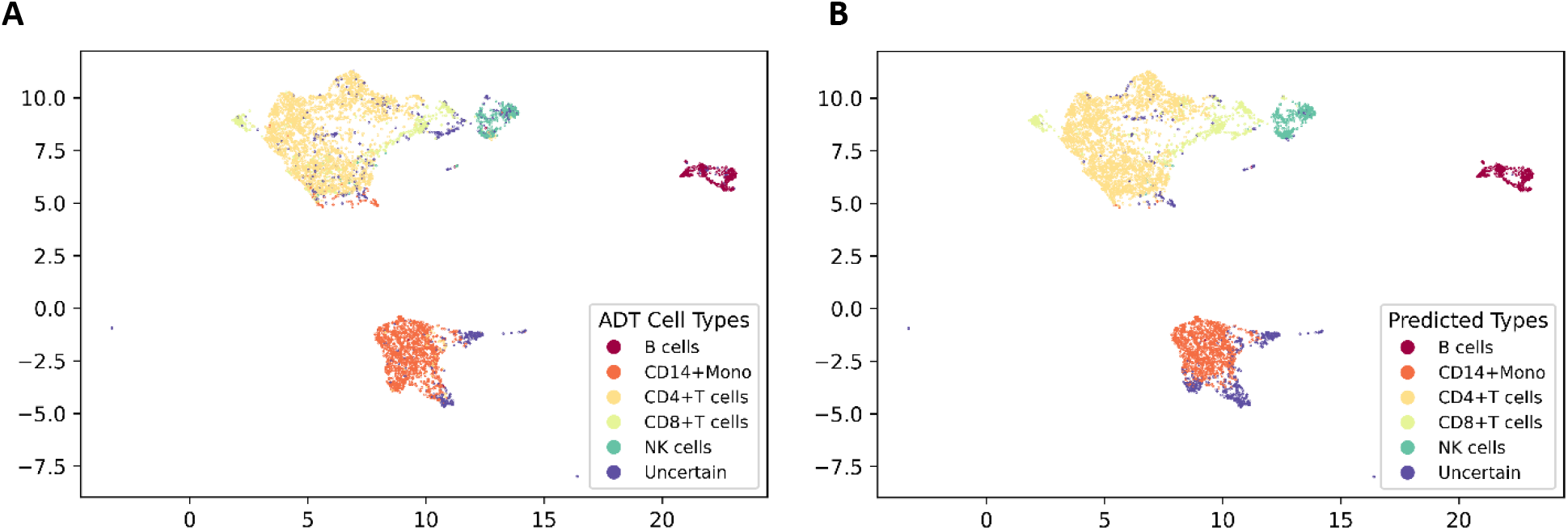
UMAP of 10X5k_PBMC RNA data on latent space. 5A is colored by confident cell types identified with ADT data. 5B is colored by the predicted confident cell types by SECANT.

### Confident cell type prediction for an in-house CITE-seq dataset

To further investigate the performance of confident cell type prediction of SECANT, we generated an in-house CITE-seq dataset of PBMCs from a healthy donor in the study of Inflammatory Bowel Disease. Cells were stained with Totalseq-A from BioLegend and are prepared using the 10x Genomics platform with Gel Bead Kit V2. The prepared assay is subsequently sequenced on an Illumina Hiseq with a depth of 50K reads per cell. Cells in this dataset are measured for their surface marker abundance through CITE-seq (3). 42 surface markers are measured for every cell, including important makers for cell type identification, such as CD3, CD4, CD8, CD14, CD16, CD19, CD56 and CD127.

Similar to joint analysis of public PBMC data, in this session, we jointly analyze public 10X10K_PBMC and our in-house data, but we only input the RNA data from our in-house CITE-seq data. Although two data are extracted from different resources, we assume the majority of immune cell types are similar in two data, thus we want to investigate if SECANT can predict confident cell types for our in-house data by borrowing ADT information from the public PBMC dataset. In Figure 6A, we show the prediction results of SECANT on a UMAP plot constructed with our in-house RNA data. Due to the experimental design that a low concentration of antibodies has been used, the ADT level remains low for most surface markers, e.g. CD3, thus we fail to build confident cell types through gating. Although we cannot compare our results with a good standard, we plot the protein abundance on UMAP to partially validate our results. As shown in Figure 6B, although it is difficult to identify cell types through CD3, we can verify B cells with the abundance of CD19 marker, CD4+ T cells with CD4 marker, CD8+ T cells with CD8 marker, Monocytes with CD14 and CD16 markers, and NK cells with CD56 and CD127 markers. Overall, the prediction results from SECANT are highly supported by the abundance of important cell surface markers. To further assess the prediction results, we compared our results with the prediction results from SingleR, a popular automatic annotation method for scRNA-seq data based on reference datasets (Aran et al. 2019), which are shown in Supplemental Figure 9. In general, cells predicted as B cells, T cells, NK cells and monocytes from SECANT and SingleR are highly consistent. Cells labeled as “uncertain” cell type by SECANT are mixture of dendritic cells, macrophages and other cell types, which is as expected since those cell types are not classified as confident cell types in the input ADT label. A confusion matrix showing the concordance of predication results from SECANT and SingleR is summarized in Supplemental Table 3, which is generally consistent with our conclusions from UMAP plot. Compared to SingleR, our proposed SECANT can provide additional annotation for cell types that are difficult to be identified with RNA markers, e.g. CD4+ T cells and CD8+ T cells, but readily to be classified with ADT markers.

**Figure 6.**
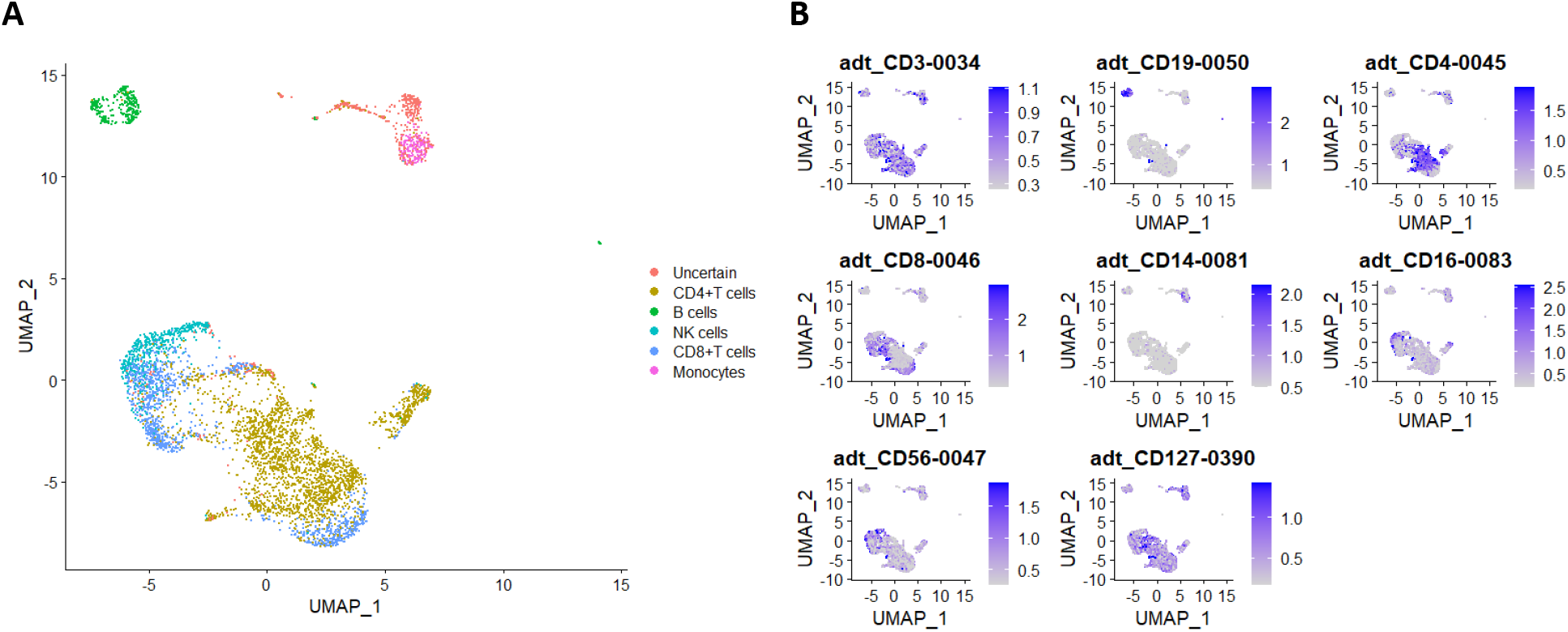
Confident cell type prediction for in-house PBMC dataset, visualized by UMAP plot constructed with RNA data from in-house CITE-seq data. 6A is colored by predicted confident cell types by SECANT. 6B is colored by surface protein abundance for some important CD markers.

## Discussion

In this study, we have developed SECANT, a biology-guided semi-supervised method for cell clustering, cell type classification and annotation for analyzing CITE-seq alone or with scRNA-seq data. Different from other existing tools for single cell multi omics, SECANT utilizes a biology-driven approach and considers that cell surface protein data can provide confident cell type labels, which are assumed to be gold standard in single cell proteomics experiments such as flow cytometry and mass cytometry, and thus should be used to guide cell clustering with RNA data. Our proposed method is developed based on model-based semi-supervised learning, and we introduce a probabilistic concordance matrix to implement ADT constraints as well as for cluster annotation. When several related scRNA-seq data are available, jointly analyzing CITE-seq and scRNA-seq data with SECANT can provide annotation of confident cell types, which are constructed with ADT data from CITE-seq, for cells from scRNA-seq, and the ADT-guided clustering performance is expected to enhance.

Still, several limitations exist for SECANT. First, in this study, the input of SECANT from ADT data is the confident cell type label built through manual gating, which undergoes a relatively subjective process. For example, a less conservative gating approach will introduce noise to cell label, while a more conservative approach will lead to the loss of sample size. In practice, as a preliminary step of SECANT, we recommend researchers to gate cells more conservatively, and leave cells on the boundary as “uncertain” cell type (e.g., Supplemental Fig. 2). SECANT is designed to fully utilize cells with uncertain cell type identified with ADT data, thus a conservative gating approach wouldn’t lead to the loss of sample size, but could sufficiently reduce the labeling noise. In addition, other methods, e.g., auto gating (Lian et al. 2020), can also be used to build confident cell type label as the input for SECANT. Second, SECANT employs stochastic gradient descent (SGD) for optimization, which is a computationally expensive approach. To speed up, we implement our algorithm in PyTorch (a Python library from Facebook) and utilizes vectorization. Although PyTorch is well-known for deep learning, it can be used to optimize a target function through SGD without building neural networks. Also, it is extremely convenient with PyTorch to use graphics processing units (GPU) for strong acceleration, and we have implemented our algorithm with GPU setting (e.g., freely available Google Colab). Third, to estimate the configuration of the matrix form of the concordance matrix *C* without any prior knowledge, currently we need to run all possible configurations, and then select the one with the maximum log-likelihood. To speed up, one can run our method in parallel, each thread running a different configuration. In addition, we are working on to develop an alternative approach for optimization, which utilizes the alternating direction method of multipliers (ADMM), in the future study. The alternative approach is expected to be more efficient than the current approach.

In summary, we propose a novel statistical method, SECANT, which utilizes model-based semi-supervised learning for surface protein guided cell clustering, classification and annotation with CITE-seq data or joint analysis with CITE-seq and scRNA-seq data. Our model framework can be extended to accommodate single cell data from more than two data sources, or to analyze data from other study fields. Additionally, our well-designed in-house CITE-seq dataset will be valuable for researchers to develop novel methods. We believe SECANT would quickly gain popularity among medical researchers, particularly in immunology filed.

## Methods

### Statistical models

We first denote *L_i_* the confident cell type label for cell *i* obtained from ADT data. We assume there are in total *M* confident cell types identified with ADT data, and the support of *L* is {1,2,…, *M, M* + 1}, where *L* = 1,2,…, *M* corresponds to each of the confident cell types, and *L* = *M* + 1 refers to the additional “uncertain” group. We then denote *Z_i_* the cell cluster label for cell *i* estimated from RNA data. Assuming the total number of clusters is *K*, the support of *Z* is {1,2,…, *K*}. The core assumption of our approach is that cells shouldn’t fall into the same cluster from RNA data if they are identified as different confident cell types (not including the “uncertain” group) with ADT data. For example, if one cell is identified as a T cell and another as a B cell confidently from ADT data, then we should avoid these two cells being clustered together from RNA data. As an exception, this constraint doesn’t apply to cells falling into the “uncertain” group, which makes our assumption plausible in practice. This constraint can be described mathematically as follows:

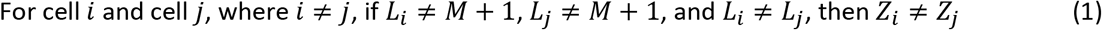

Equivalently, we can state our constraint (1) in a statistical way by introducing a concordance matrix *C*_(*m*+1)×*K*_, as shown in Table 2. We denote *p_mk_* in the matrix the conditional probability *P*(*L_i_* = *m|Z_i_* = *k*) for cell *i*. The total number of parameters in *C* is *K*. Under constraint (1) and general assumptions, we have the following constraints on the first *M* rows of *C*, denoted by *C*[1: *M*,:], of which each row corresponds to a confident cell type:

1. Each column of *C*[1: *M*,:] contains exactly one parameter, which ranges from 0 to 1;
2. Each row of *C*[1: *M*,:] contains at least one parameter, which ranges from 0 to 1;
3. All entries of *C*[1: *M*,:] except for the *K* parameters are 0

**Table 2.** An example of concordance matrix with *M* confident cell types identified with ADT data and *K* clusters identified with RNA data under ADT guidance. Each entry of the matrix represents the conditional probability of a cell belong to a certain cell type given its clustering category.

|  |  | Clusters from RNA data |  |  |  |  |  |  |
| --- | --- | --- | --- | --- | --- | --- | --- | --- |
| | | Cluster 1 | Cluster 2 | ... | Cluster $k$ | Cluster $k+1$ | ... | Cluster $K$ |
| Confident cell types from ADT data | Cell Type 1 | $p_{11}$ | 0 | | 0 | 0 | | 0 |
| | Cell Type 2 | 0 | $p_{22}$ | | 0 | 0 | | 0 |
|  | ... |  |  |  |  |  |  |  |
| | Cell Type $m$ | 0 | 0 | | $p_{mk}$ | $p_{m,k+1}$ | | 0 |
|  | ... |  |  |  |  |  |  |  |
| | Cell Type $M$ | 0 | 0 | | 0 | 0 | | $p_{MK}$ |
| | Uncertain* | $1 - p_{11}$ | $1 - p_{22}$ | | $1 - p_{mk}$ | $1 - p_{m,k+1}$ | | $1 - p_{MK}$ |

The last row of *C*, referring to the “uncertain” group, can then be decided with parameters implemented in *C*[1: *M*,:]. In general, there are multiple configurations of the matrix form of *C* that fulfill the constraints described above, and there are *K* parameters to be estimated for each configuration. We will discuss more about the concordance matrix *C* in the next session. After introducing the concordance matrix *C*, we can then write out the likelihood function.

### Scenario 1: CITE-seq data only

We denote *Y* the latent space of RNA data, and each element *Y_ij_* represents the value for feature *j* in cell *i*, where *i* runs from 1 to the total number of cells *N*, and *j* runs from 1 to the number of dimensions in latent space *D*. We further assume the total number of clusters identified from RNA data is *K*. The likelihood can be written as:

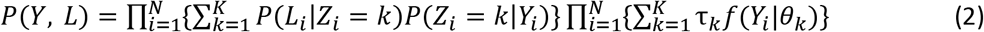

where 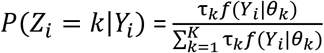 refers to the posterior probability of cell *i* belonging to cluster *k*, and *τ_k_* = *P*(*Z_i_* = *k*) refers to the proportion of cluster *k. f*(*Y_i_|θ_k_*) refers to the cluster-specific distribution of RNA data on latent space. In this study, we assume 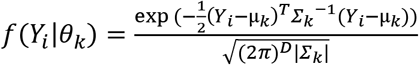, the probability density function (pdf) of multivariate Gaussian distribution with *θ_k_* = {μ_*k*_, *∑_k_*}, where μ_*k*_ stands for a *D*-dimensional cluster-specific mean vector and *∑_k_* stands for a *D* by *D* cluster-specific covariance matrix. In addition, *P*(*L_i_|Z_i_* = *k*) are elements of the concordance matrix *C* we mentioned before.

### Scenario 2: CITE-seq data and scRNA-seq data

We denote *Y*^(1)^ the latent space of RNA data from CITE-seq, and each element 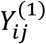 represents the value for feature *j* in cell *i*, where *i* runs from 1 to *N*_1_, and j runs from 1 to *D*. We then denote *Y*^(2)^ the latent space of RNA data from scRNA-seq, and each element 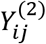 represents the value for feature *j* in cell *i*, where *i* runs from *N*_1_ + *1* to *N*_1_ + *N*_2_, and *j* runs from 1 to *D*. We assume *Y*^(1)^ and *Y*^(2)^ have batch effect corrected, and the features are exactly matched. We denote *L*^(1)^ the ADT confident cell types obtained from CITE-seq data. We further assume the common total number of clusters identified from RNA data is *K*. Similar to Scenario 1, the likelihood can be written as:

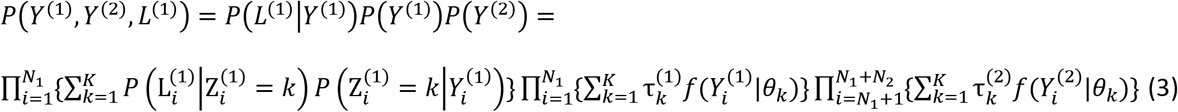

where 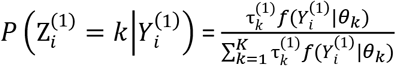 refers to the posterior probability of cell *i* belonging to cluster *k*, and 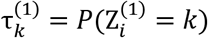 refers to the proportion of cluster *k* in RNA data from CITE-seq. Similarly, 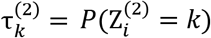 refers to the proportion of cluster *k* in RNA data from scRNA-seq. 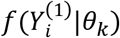 and 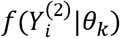 refer to the cluster-specific distribution of RNA data from CITE-seq and scRNA-seq on latent space, respectively. Similar to Scenario 1, we assume 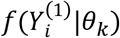 and 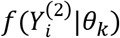 are the pdf of multivariate Gaussian distribution with *θ_k_* = {μ_*k*_, *∑_k_*}. For model flexibility, data-specific cluster proportions, 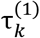 and 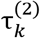, are allowed to differ. Note that the two data sources (after batch effect correction) share the common cluster-specific parameters. Again, *P*(*L_i_*|*Z_i_* = *k*) are elements of the concordance matrix *C*, and we assume the concordance matrix is common between two data sources. For prediction of confident cell types for scRNA-seq data, denoted by *L*^(2)^, we can compute the posterior probability of cell *i* belonging to confident cell type *m*, 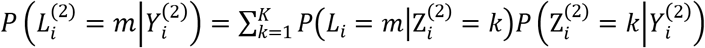, where 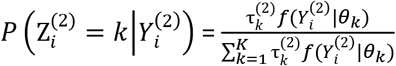 refers to the posterior probability of cell *i* belonging to cluster *k* for scRNA-seq data.

### Modeling and space reduction of concordance matrix *C*

The concordance matrix *C* we proposed in our model has two major functions: 1) to associate ADT data and RNA data by considering the confident cell types label from ADT data as guidance for cell clustering with RNA data; 2) to provide the confident cell type annotation for each cluster. In general, a concordance matrix with *M* confident cell types and *K* clusters, as shown in Table 2, has 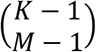 different configurations of matrix form that fulfill the constraints we described above. Here the notation 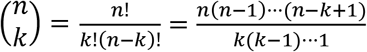 refers to the number of *k*-combinations of a set *S* with *n* elements. The number 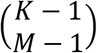 is derived from an analogy of listing all possible configurations for arranging *K* balls into *M* different boxes providing that each box has at least one ball. For example, when *K* = 11 and *M* = 5, the total number of different configurations is 210. In practice, one can specify a couple of plausible configurations with prior knowledges, or to run our algorithm with all possible configurations in parallel, and then select the final configuration with the largest log-likelihood (Supplemental Fig. 10). Although the number 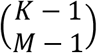 can be large, a great number of matrix forms are actually not practical at all, for instance, when one confident cell type is assumed to correspond to six cell clusters and the other five confident cell types each corresponds to only one cell cluster. As a result, the number can be largely reduced by restricting the maximum number of clusters a confident cell type corresponds to, denoted by 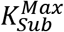. For example, when we set 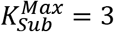, the total number of matrix forms is reduced to 45 from 210 for the situation when *K* =11 and *M* = 5. 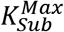 can be selected based on prior biological knowledge, or information from a UMAP plot (e.g., Fig. 4A).

### Optimization method

We use stochastic gradient descent (SGD) method to directly optimize the log-likelihood (by minimizing the negative log-likelihood) of observed data, where the likelihood function for each scenario (1. CITE-seq data only; 2. CITE-seq data and scRNA-seq data) is defined above. The parameters to be estimated through SGD include cluster-specific parameters {*τ_k_*, μ_*k*_, *∑_k_*} from the clustering part, and the parameter *p_mk_’s* in the concordance matrix *C*. As described in the previous session, each configuration of the matrix form of *C* is also a parameter in the likelihood function, which can be maximized through parallel computing (each thread with a different configuration). One of the computational advantages of our method is that our algorithm is implemented with PyTorch and users can run it on computers with GPU accelerations (e.g. freely available Google Colab).

### Initialization of algorithm

For the initialization of clustering related parameters {*τ_k_*, μ_*k*_, *∑_k_*}, we exclude cells with uncertain ADT label and run separate multivariate GMM with cells from each confident cell type identified with ADT data.

The number of mixtures for each multivariate GMM is decided based on the concordance matrix *C*. For the initialization of parameters in *C*, by default we set *p_mk_* = 0.5 as the initial value. In practice, we recommend running our method with five different initializations and select the one with the largest log-likelihood. The pair-wise ARI and AMI among clustering results with different initialization for 10X10K_PBMC dataset is summarized in Supplemental Table 4, from which we observe that the clustering results are very similar.

## Software availability

Our algorithm is implemented in Python based on PyTorch, and is available on GitHub (https://github.com/tarot0410/SECANT) with a detailed tutorial.

## Data access

The public datasets we used in this study can be downloaded from 10X Genomics website (10k: https://support.10xgenomics.com/single-cell-geneexpression/datasets/3.0.0/pbmc_10k_protein_v3; 5k: https://support.10xgenomics.com/single-cell-gene-expression/datasets/3.0.2/5k_pbmc_v3_nextgem). The in-house CITE-seq dataset will be uploaded to Gene Expression Omnibus (GEO).

## Acknowledgments

This work was funded by National Institutes of Health grants [P01AI106684 to W.C., U01DK062420 to W.C., R.D.], and was also supported in part by Children’s Hospital of Pittsburgh of the UPMC Health System, and the University of Pittsburgh Center for Research Computing through the resources provided.

