## Supplementary data for "SECANT: a biology-guided semi-supervised method for clustering, classification, and annotation of single-cell multi-omics"

**Supplemental Figure 1:** General workflow illustrating how to obtain confident cell type label (for PBMCs) through manual gating with ADT data.

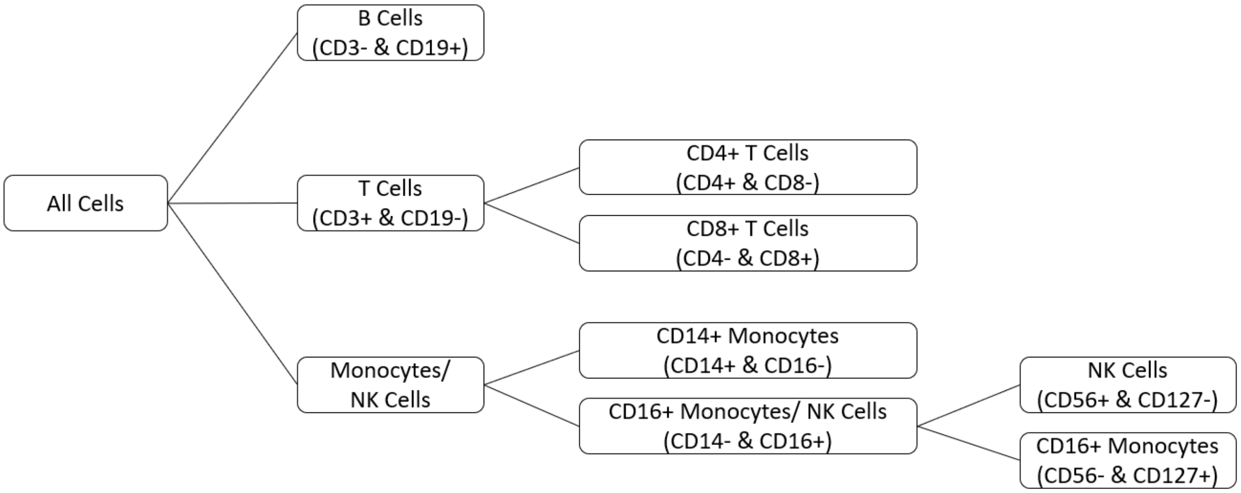

**Supplemental Figure 2:** Scatter plot of cells illustrating manual gaiting with public human 10X10k_PBMC dataset.

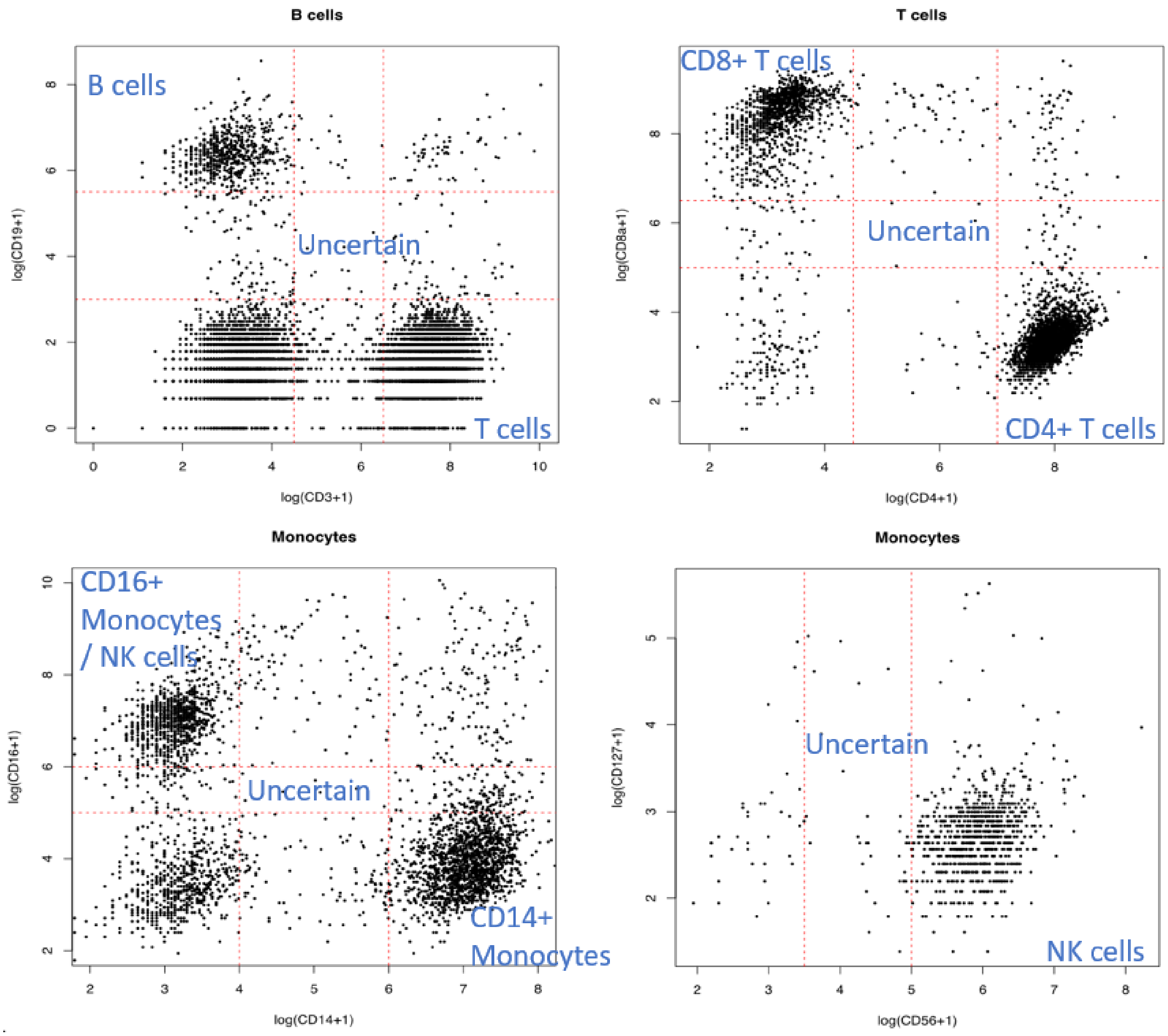


.

**Supplemental Figure 3:** Batch effect correction for RNA data from 10X10k PBMC and 10X5k PBMC CITE-seq datasets. 3A shows the UMAP of two datasets pooled together without batch effect correction. 3B shows the UMAP on latent space of two datasets after using scVI for dimension reduction and batch effect correction.

**A**

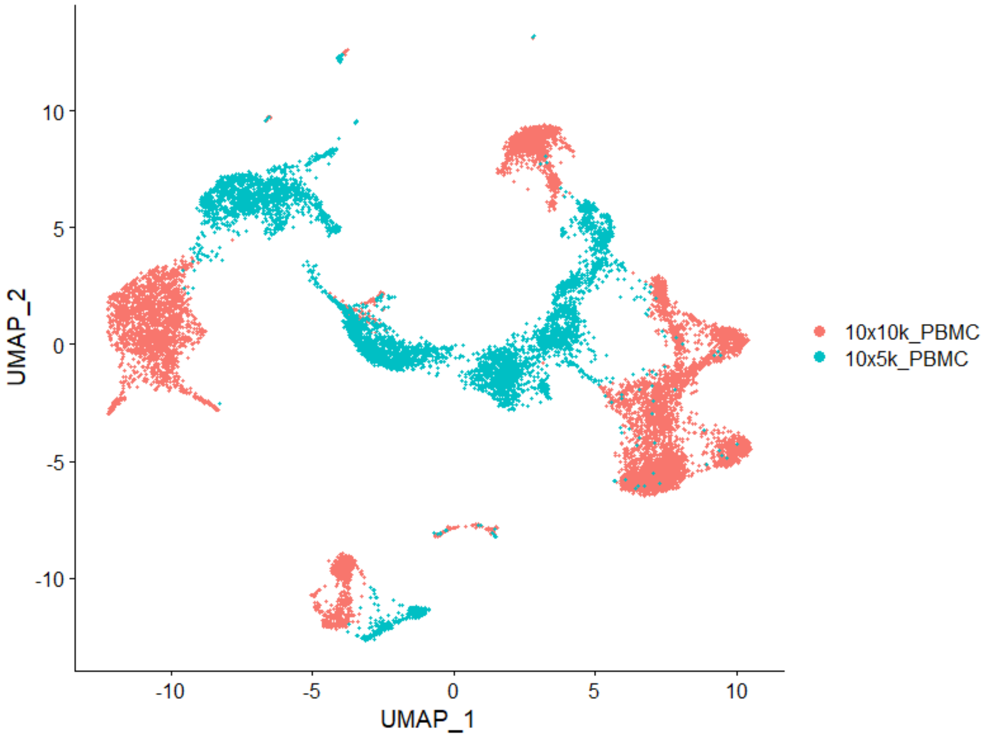


**B**

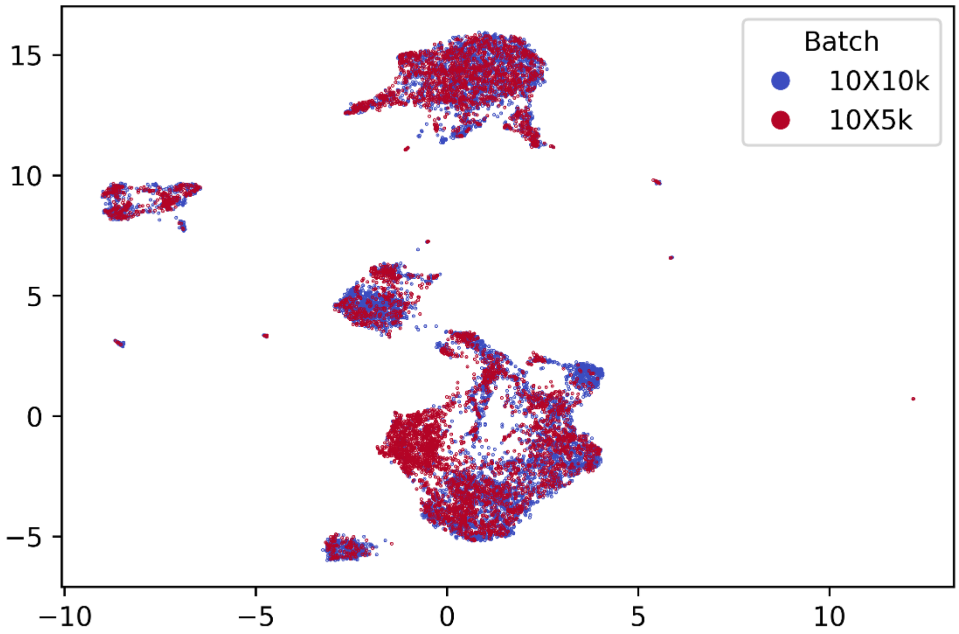

**Supplemental Figure 4:** Additional results from simulation studies. 4A shows the distribution of AMI of SECANT (with different $p^U$ setting), K-means and GMM across different sample size $N$. 4B shows the distribution of AMI of SECANT using one dataset input vs. two datasets input under various $p^U$ and $N$ settings. 4C shows the trend of proportion of cells being predicted as "uncertain" cell type with various $p^U$ and $N$ settings.

**A**

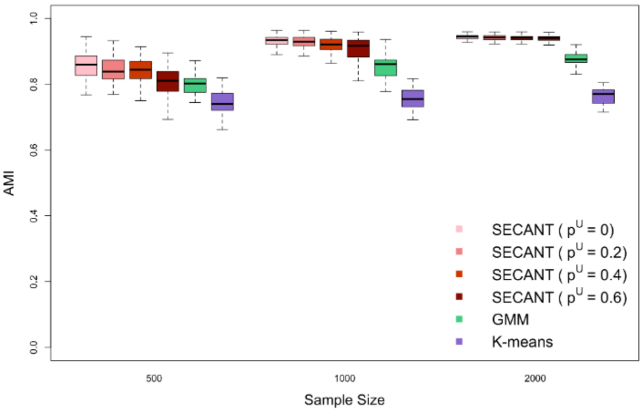


**B**

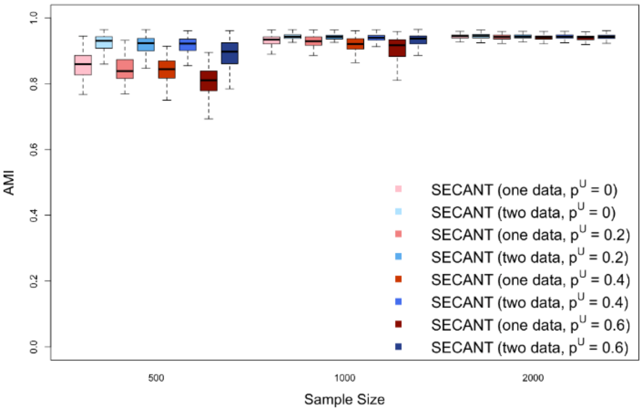


**C**

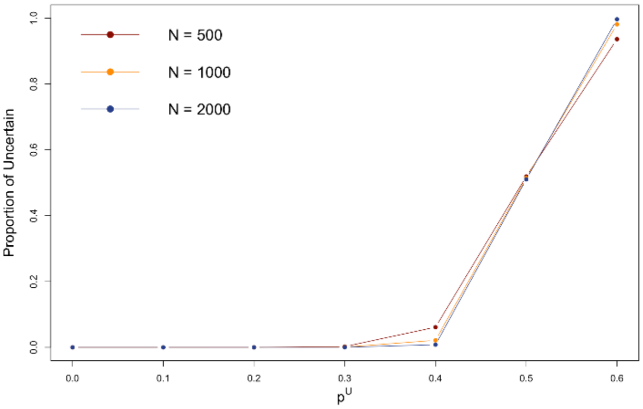

**Supplemental Figure 5:** Simulation results for the scenario of detecting novel cell cluster. 5A shows the simulated cell type label, where a cluster of cells are labeled as “uncertain”. 5B shows the ADT-guided clustering results from SECANT.

**A**

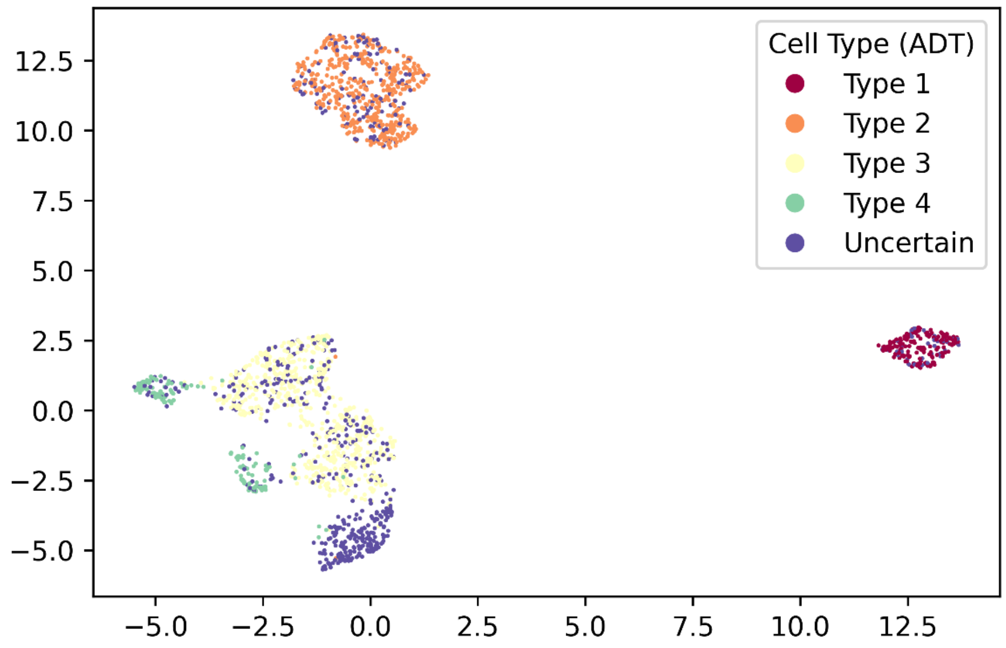


**B**

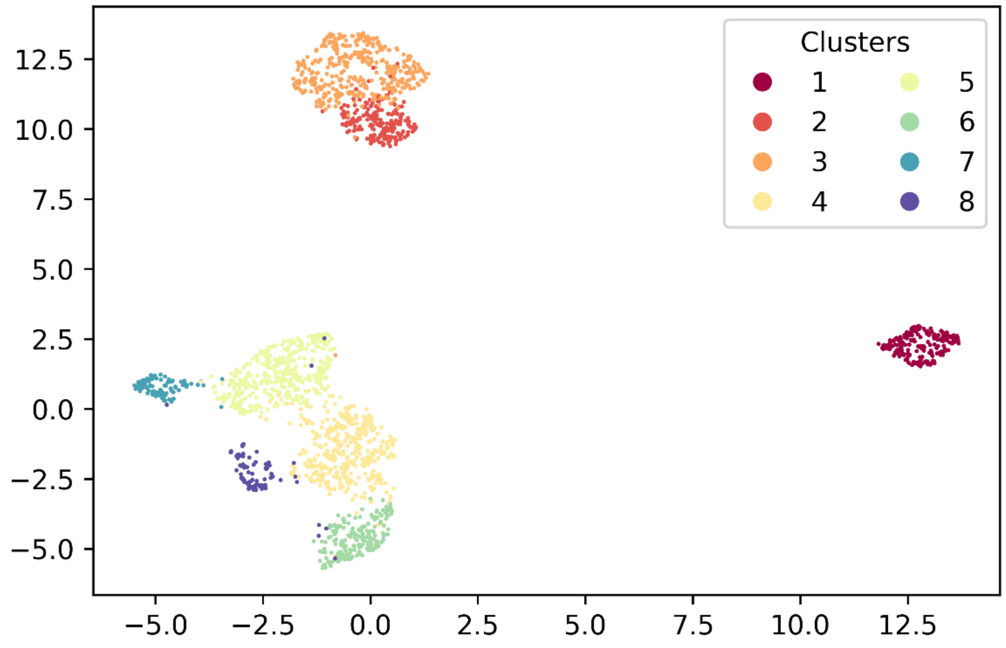

**Supplemental Figure 6:** UMAP of 10X10k_PBMC RNA data on latent space. 6A, 6B and 6C are colored by cell types identified with ADT data, clustering results from GMM, and results from Seurat, respectively.

**A**

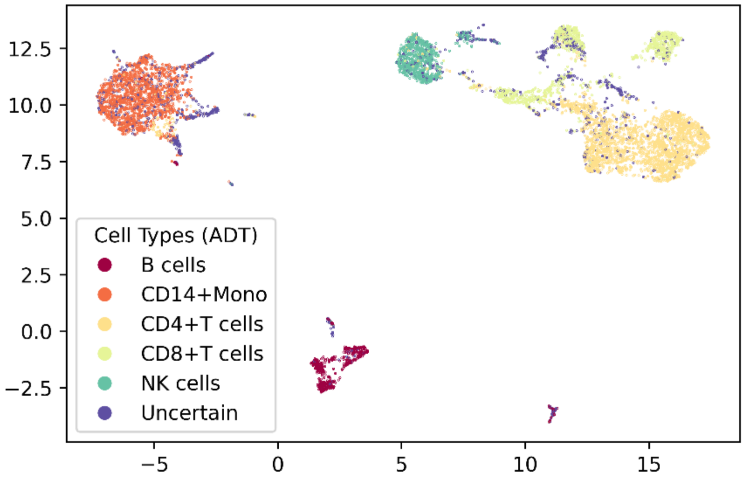


**B**

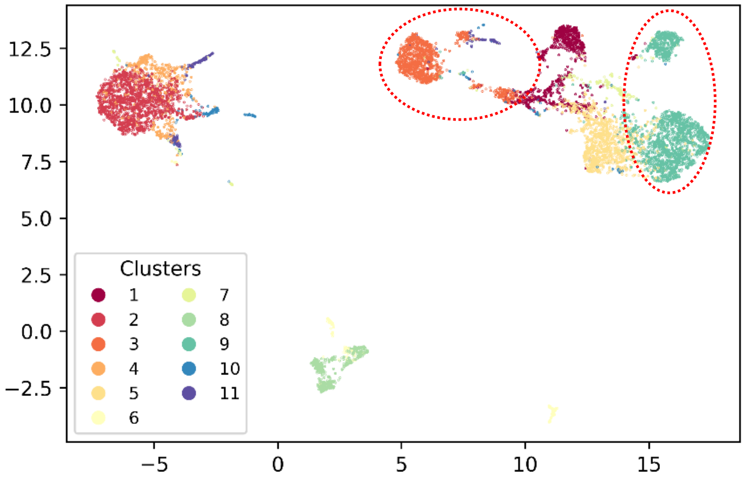


**C**

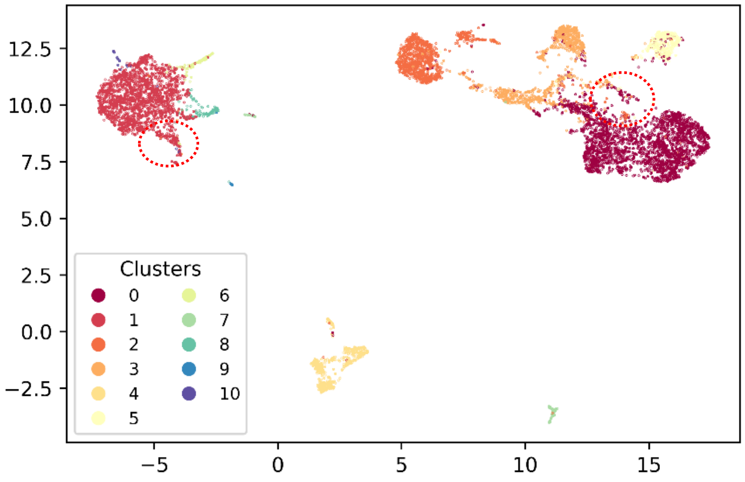

**Supplemental Figure 7:** UMAP of 10X5k_PBMC RNA data on latent space. 7A, 7B, 7C and 7D are colored by cell types identified with ADT data, clustering results from SECANT, clustering results from GMM, and clustering results from Seurat, respectively.

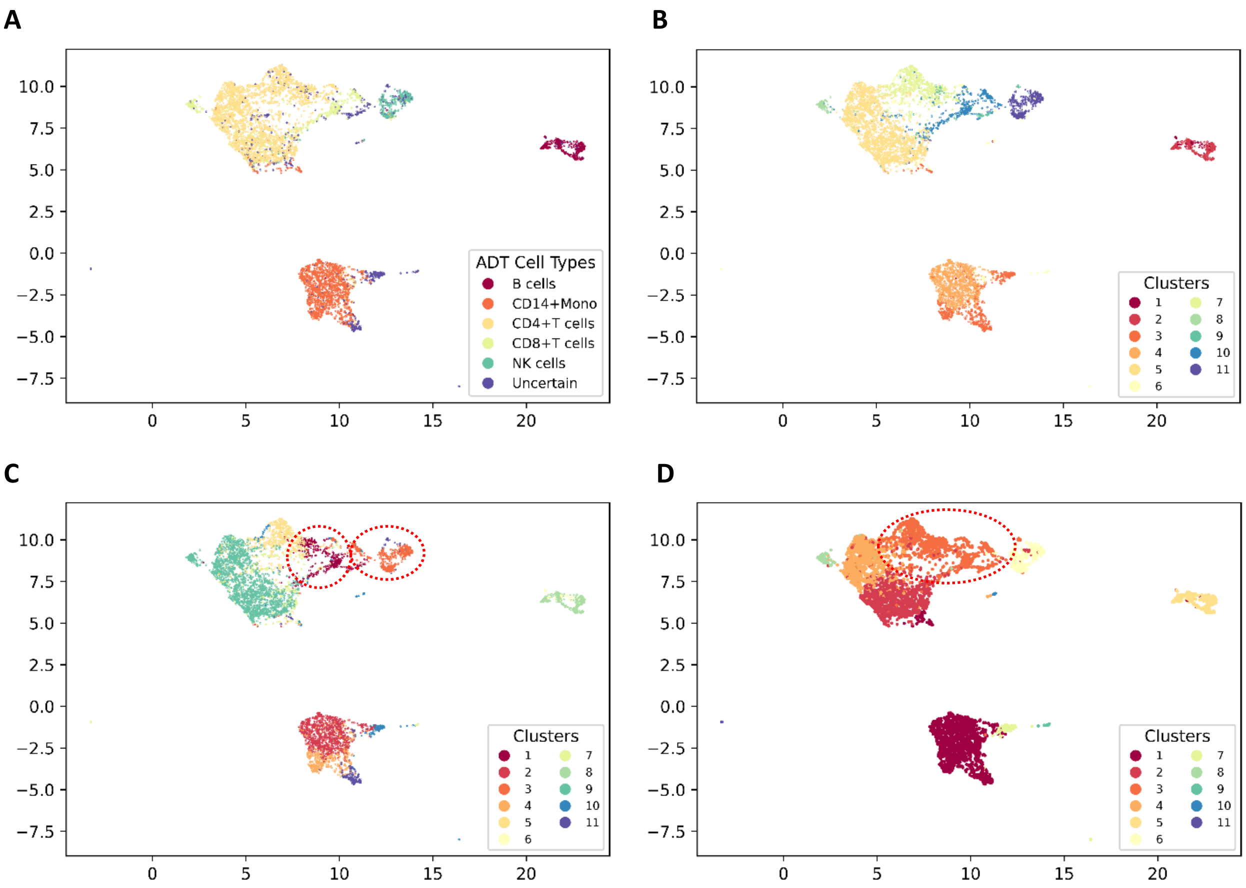

**Supplemental Figure 8:** Boxplot showing the distribution of maximum posterior probability for cells with correctly predicted labels (Group 1) vs cells with incorrectly predicted labels (Group 2).

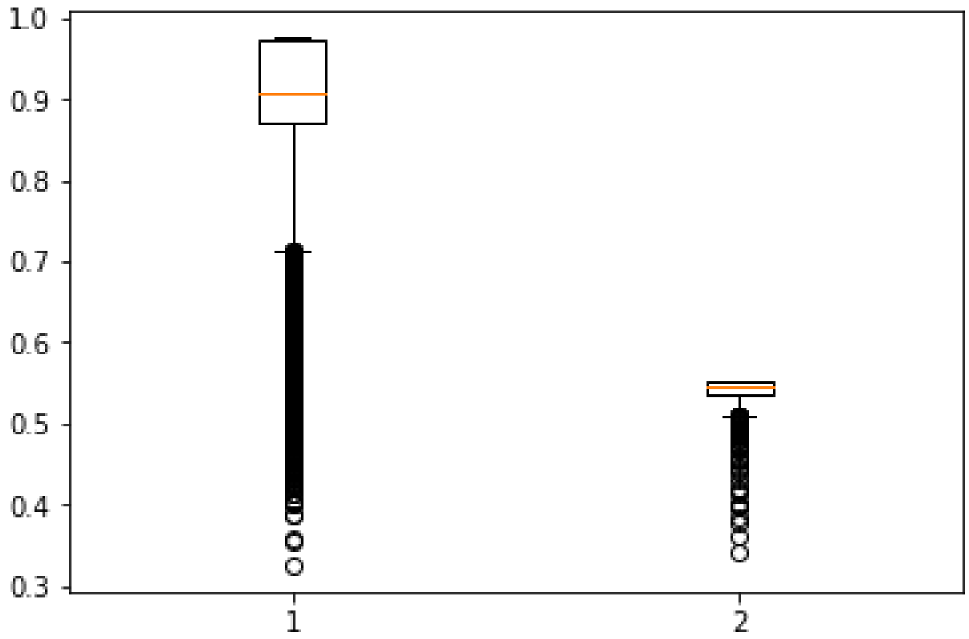

**Supplemental Figure 9:** Cell type prediction for in-house PBMC dataset, visualized by UMAP plot constructed with RNA data from in-house CITE-seq data. 9A is colored by predicted confident cell types by SECANT. 9B is colored by predicted cell types by SingleR.

**A**

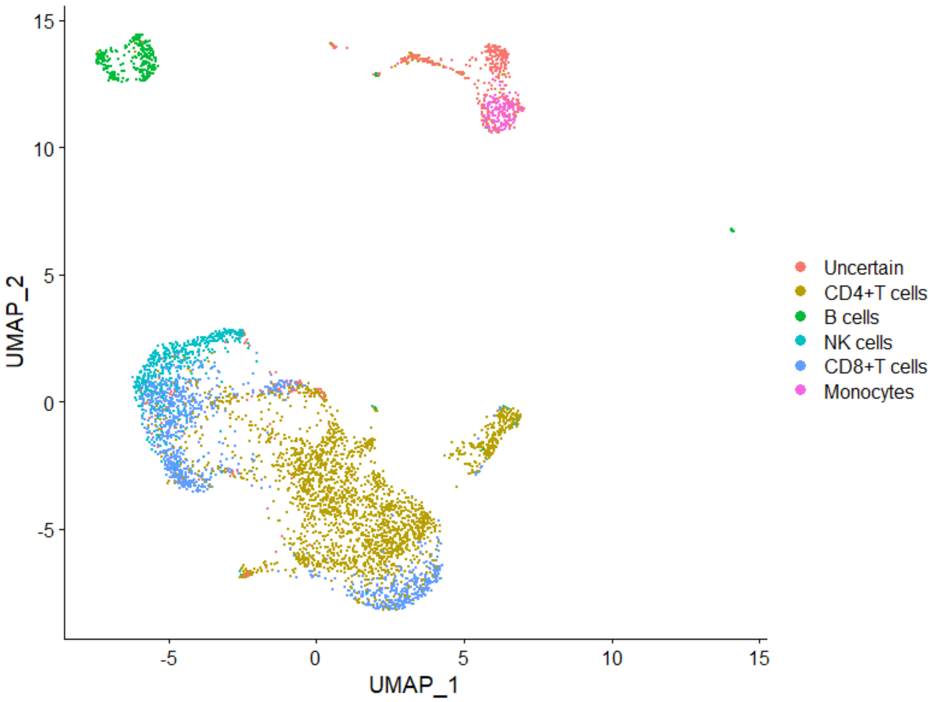


**B**

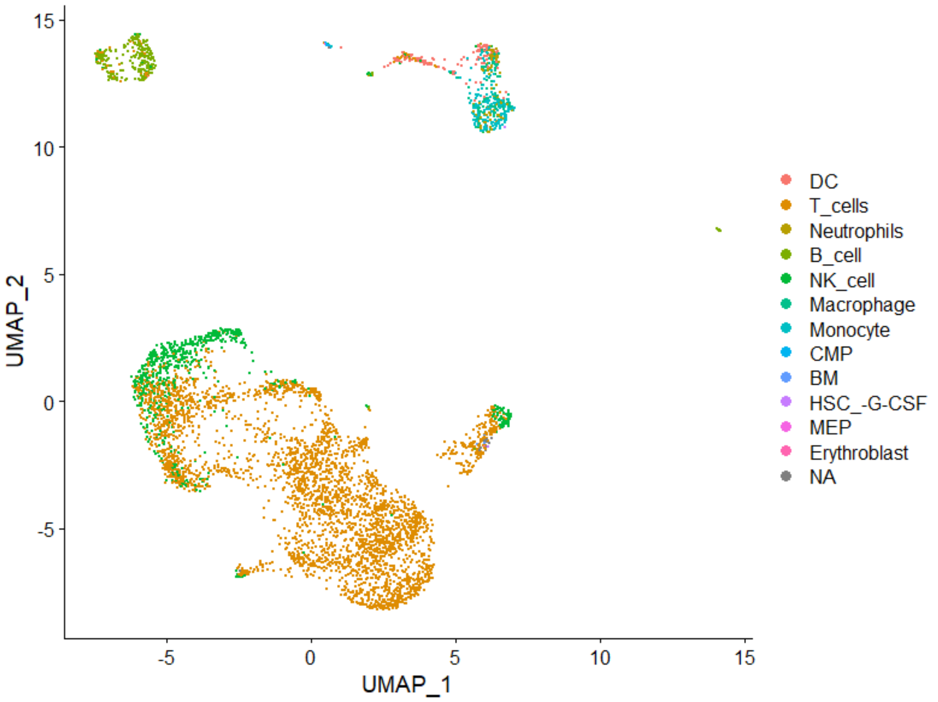

**Supplemental Figure 10:** Distribution of log-likelihood for each different matrix form of concordance matrix. Five different initializations are applied for each matrix form.

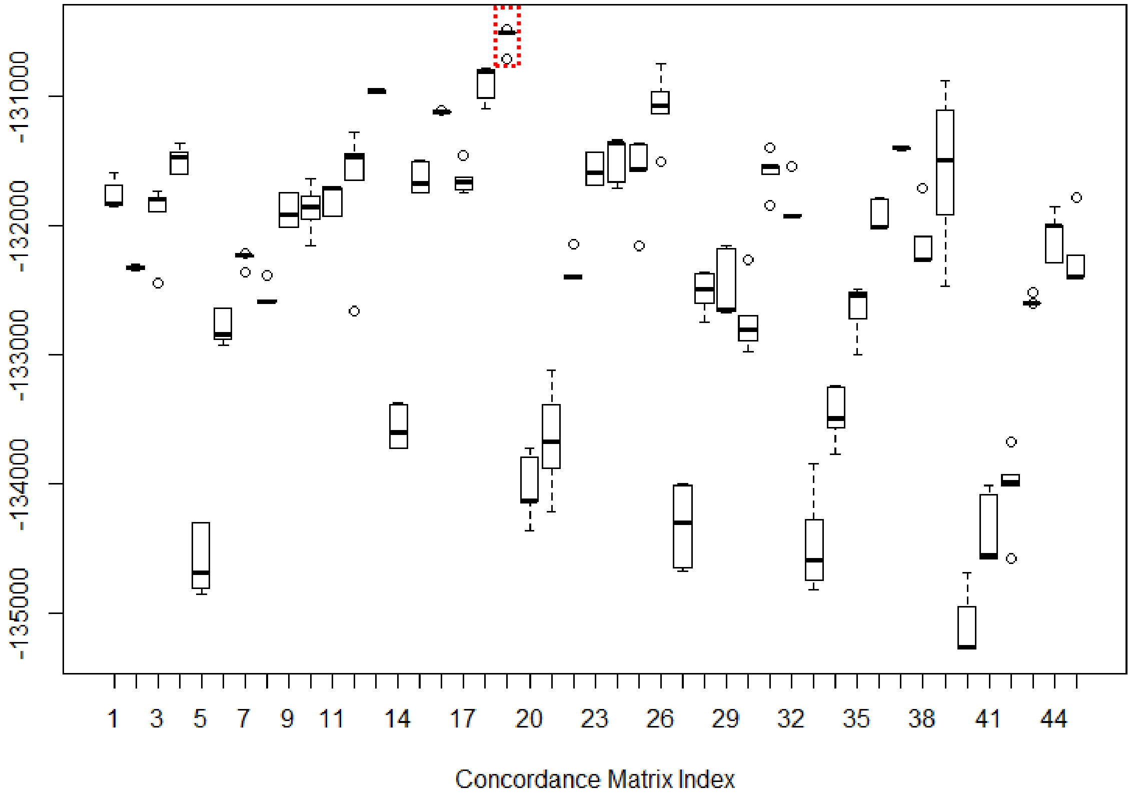

**Supplemental Table 1:** Estimated concordance matrix under the simulation scenario of detecting novel cell cluster.

| | Cluster 1 | Cluster 2 | Cluster 3 | Cluster 4 | Cluster 5 | Cluster 6 | Cluster 7 | Cluster 8 |
|---|---|---|---|---|---|---|---|---|
| Type 1 | 0.82 | 0 | 0 | 0 | 0 | 0 | 0 | 0 |
| Type 2 | 0 | 0.809 | 0.79 | 0 | 0 | 0 | 0 | 0 |
| Type 3 | 0 | 0 | 0 | 0.776 | 0.764 | 0 | 0 | 0 |
| Type 4 | 0 | 0 | 0 | 0 | 0 | 0 | 0.772 | 0.785 |
| Uncertain | 0.18 | 0.191 | 0.21 | 0.224 | 0.236 | 1 | 0.228 | 0.215 |

**Supplemental Table 2:** Confusion matrix of predicted confident cell types vs. actual cell types built from ADT data.

| | **Actual** | | | | | |
|---|---|---|---|---|---|---|
| **Predicted** | B cells | CD14+ Monocytes | CD4+ T cells | CD8+ T cells | NK cells | Uncertain |
| B cells | 327 | 0 | 0 | 0 | 0 | 35 |
| CD14+ Monocytes | 0 | 1052 | 13 | 0 | 0 | 29 |
| CD4+ T cells | 0 | 34 | 2271 | 92 | 3 | 108 |
| CD8+ T cells | 0 | 0 | 84 | 365 | 2 | 131 |
| NK cells | 1 | 0 | 14 | 5 | 278 | 55 |
| Uncertain | 1 | 220 | 105 | 9 | 14 | 279 |

**Supplemental Table 3:** Confusion matrix of predicted general cell types from SECANT vs. SingleR.

| | SECANT | | | | | |
|---|---|---|---|---|---|---|
| **SingleR** | B cells | CD4+ T cells | CD8+ T cells | Monocytes | NK cells | Uncertain |
| B_cell | 256 | 0 | 0 | 0 | 0 | 4 |
| BM | 0 | 3 | 0 | 0 | 0 | 0 |
| CMP | 0 | 2 | 0 | 0 | 0 | 16 |
| DC | 2 | 16 | 0 | 2 | 0 | 105 |
| Erythroblast | 1 | 1 | 0 | 0 | 0 | 0 |
| HSC_-G-CSF | 1 | 2 | 0 | 0 | 0 | 1 |
| Macrophage | 327 | 0 | 0 | 35 | 0 | 73 |
| MEP | 0 | 2 | 0 | 0 | 0 | 0 |
| Monocyte | 0 | 3 | 0 | 95 | 0 | 80 |
| Neutrophils | 0 | 0 | 0 | 25 | 0 | 54 |
| NK_cell | 17 | 82 | 151 | 0 | 407 | 49 |
| T_cells | 33 | 2267 | 959 | 1 | 97 | 100 |

**Supplemental Table 4:** Pair-wise ARI and AMI among clustering results with different initialization.

ARI:

| | Initial 1 | Initial 2 | Initial 3 | Initial 4 | Initial 5 |
|---|---|---|---|---|---|
| Initial 1 | 1.0000 | 0.9996 | 0.9892 | 0.9828 | 1.0000 |
| Initial 2 | 0.9996 | 1.0000 | 0.9887 | 0.9833 | 0.9996 |
| Initial 3 | 0.9892 | 0.9887 | 1.0000 | 0.9795 | 0.9892 |
| Initial 4 | 0.9828 | 0.9833 | 0.9795 | 1.0000 | 0.9828 |
| Initial 5 | 1.0000 | 0.9996 | 0.9892 | 0.9828 | 1.0000 |

AMI:

| | Initial 1 | Initial 2 | Initial 3 | Initial 4 | Initial 5 |
|---|---|---|---|---|---|
| Initial 1 | 1.0000 | 0.9987 | 0.9804 | 0.9655 | 1.0000 |
| Initial 2 | 0.9987 | 1.0000 | 0.9793 | 0.9663 | 0.9987 |
| Initial 3 | 0.9804 | 0.9793 | 1.0000 | 0.9605 | 0.9804 |
| Initial 4 | 0.9655 | 0.9663 | 0.9605 | 1.0000 | 0.9655 |
| Initial 5 | 1.0000 | 0.9987 | 0.9804 | 0.9655 | 1.0000 |
